# Assessing the efficacy of human mesenchymal stromal cells of different tissue origins in a mouse model of kidney ischaemia reperfusion injury

**DOI:** 10.64898/2026.06.19.733188

**Authors:** Katherine Trivino-Cepeda, Francesco Amadeo, David M. Hughes, Lorenzo Ressel, Marta Garcia-Finana, Vivien Hanson, Arthur Taylor, Patricia Murray, Bettina Wilm

## Abstract

Rodent models of kidney disease have been widely used to assess the efficacy, safety and mode of action of mesenchymal stromal cells (MSCs) as therapies. However, because kidney disease models, MSC type and the methods used to assess kidney injury tend to differ between research groups, it is difficult to obtain data that are sufficiently robust and reproducible to support clinical translation.

We present here for the first time a side-by-side analysis of the performance of human MSCs derived from the most commonly used tissue sources, bone marrow (BM-), adipose- (A-) and umbilical cord (UC-), in a kidney ischaemia reperfusion injury (IRI) model in mice. For each animal, we performed a comprehensive assessment of kidney function and health by longitudinal transdermal measurements of sinistrin clearance, serum biomarker levels at the experimental endpoint, and histopathological scoring of sections from left and right kidneys. Furthermore, we tracked the MSCs by bioluminescence imaging in the injured mice to determine their viability over time and their capacity for homing to the damaged kidneys.

Our results reveal that only modest if any beneficial effects of the MSC treatments were detectable on kidney function and histology, irrespective of cell type administered. Furthermore, all three MSC types were sequestered in the lungs without reaching the kidneys, and had completely disappeared within 7 days. Our data suggest that none of the MSC types has the capability to improve renal health following IRI to a meaningful extent, questioning their suitability as a clinical therapy.

**Significance Statement:** MSCs have been proposed as efficacious cell therapies in murine models of kidney disease, with potential for clinical translation. We compare efficacy of human MSCs of different tissue origins (adipose, bone marrow and umbilical cord) in a refined mouse model of renal IRI. Only modest if any beneficial effects on kidney function and histology were detectable for all three cell types, and cells did not reach the kidneys but sequestered in the lungs where they died.

## 1. INTRODUCTION

Kidney disease is highly prevalent in societies with a high incidence of obesity, diabetes and an ageing population. Chronic kidney disease (CKD) is not curable and inevitably progresses to end stage kidney disease (ESKD), the only treatments for which are dialysis and kidney transplantation. CKD can be caused by drug treatment or other medical interventions, and often initially presents as acute kidney injury (AKI). Because of the large socioeconomic burden of kidney disease and its leading role in causing mortality, there is an urgent need to develop better treatments and therapies to prevent kidney damage and slow its progression to ESKD (Kovesdy, 2022). Rodent models for AKI and CKD addressing different aspects of kidney injury and disease have been developed to investigate disease mechanisms and assess the potential of therapeutic interventions.

In the last 20 years, there has been an intensive effort to determine the efficacy and safety of cells as regenerative therapeutic agents in acute and degenerative diseases of the kidney and other organs (Borges and Schor, 2018; Fazekas and Griffin, 2020; Petrosyan et al., 2022; Shokoohmand et al., 2025; Torrico et al., 2022), with mesenchymal stromal cells (MSCs) as an attractive source (Amadeo et al., 2021; Han et al., 2025; Musial-Wysocka et al., 2019; Wang et al., 2024b). MSCs have been shown to have immunomodulatory roles via secretion of factors which contribute to tissue repair and regeneration, by homing to the area of injury (Musial-Wysocka et al., 2019; Torrico et al., 2022; Wise et al., 2014). However, their mechanisms of action are still not clear and cell tracking studies have shown that following intravenous administration, MSCs are sequestered in the lungs and do not home to the kidneys (Amadeo et al., 2023a; Amadeo et al., 2022; Fischer et al., 2009; Geng et al., 2014; Hoogduijn et al., 2013; Shokoohmand et al., 2025; Taylor et al., 2020). The most commonly used MSCs in preclinical models and clinical trials include human bone marrow- (BM-), adipose tissue- (A-) and umbilical cord-derived MSCs (UC-MSCs) which can usually be harvested in sufficient quantities and with relative ease, making them attractive candidates for therapies (Han et al., 2025). Despite this, the number of cells needed for cell therapies requires culture and expansion of the cells, which can affect the consistency of the cell characteristics and harmonisation and standardisation of their application. We have recently reported that harmonised culture and standardised potency assays of BM-MSCs, A-MSCs and UC-MSCs between three laboratories as part of the RenalToolBox EU ITN Network [https://www.renaltoolbox.org], resulted in comparable outcomes for each cell type (Calcat-i-Cervera et al., 2023). Despite the similarities between the cells, using a range of in vitro assays, we found that conditioned medium derived from BM-MSCs had a significantly enhancing effect on endothelial tubulogenesis and migration, while conditioned medium from A-MSCs elicited a stronger immune-suppressive response in immunomodulatory experiments. On the other hand, conditioned medium from UC-MSCs had the least potency in all assays tested.

Previously, MSCs from different tissue sources have been used interchangeably in different rodent strains, utilising different kidney injury models and different measures of kidney function, making it very difficult to provide a clear statement regarding efficacy, mode of action or suitability of one cell type over the other (Cai et al., 2014; Cao et al., 2010; da Silva et al., 2015; Fang et al., 2012; Geng et al., 2014; Hauser et al., 2010; Jang et al., 2014; Morigi et al., 2008; Shih et al., 2013; Wise et al., 2014). These and other studies used kidney injury models based on folic acid, rhabdomyolysis or ischaemia reperfusion injury (IRI) in mice or rats, transplanted BM-, UC- or A-MSCs and used venous or arterial routes of administration. Interestingly, these studies reported that the MSCs tested had renoprotective effects determined by decreased levels of serum creatinine and/or blood urea nitrogen, and showed improved tissue integrity in histopathological assessments, including the status of tubular cells and necrosis. Despite these outcomes, it is important to note that especially in mouse studies, longitudinal assessments for serum biomarkers are very difficult to perform due to the fact that blood cannot be collected repeatedly without impairing animal welfare (Meyer et al., 2020; NC3Rs). Assessments of kidney health and function either take place at the experimental end point or with several cohorts of mice that are terminated at different time points of the study, therefore lacking information on individual trajectories, and making it problematic to draw conclusions of the effect of cell therapies over the entire experimental time window and across animal groups. In addition, the published studies lack harmonised experimental design regarding animal species, injury model, cell type, dosing and route of administration, making interpretation of efficacy difficult.

We have previously refined an ischaemia reperfusion injury (IRI) mouse model of AKI in which after a standardised pre-surgical anaesthetic time of 30 mins, the pedicles of both kidneys were clamped simultaneously for 27.5 minutes (Harwood et al., 2022). We assessed the injury response by including the longitudinal measurements of FITC-sinistrin renal clearance using a transdermal device which is compliant with the principles of the 3Rs (Scarfe et al., 2018a). In this assessment, FITC-sinistrin is exclusively filtered by the glomeruli and used to measure the glomerular filtration rate (GFR). This approach allows the clinically relevant classification of renal function using the RIFLE criteria which had originally been defined by the ADQI group (Bellomo et al., 2004). The RIFLE criteria utilise percentage reduction of GFR and can be adopted in preclinical rodent models by classifying kidney disease into risk, injury, failure, loss and end stage kidney disease (RIFLE). The minimally-invasive measurements of renal function make this experimental approach suitable for longitudinal assessments of the efficacy of cell therapies.

An additional aspect associated with administering MSCs as therapies in rodent models, is the question of safety and long-term fate of the cells. Intravenous administration of MSCs is a common delivery route in rodent studies; however, it has been reported to lead to sequestration and death of MSCs in the lung, also known as the pulmonary first pass effect (de Witte et al., 2018; Eggenhofer et al., 2012; Fischer et al., 2009; Leibacher and Henschler, 2016; Patrick et al., 2020; Scarfe et al., 2018b). This is in contrast to reports that the presence of an injury can lead to the migration of a proportion of the intravenously administered MSC population toward the site of kidney damage (Burks et al., 2015; Jang et al., 2014; Schubert et al., 2018; Wise et al., 2014; Xing et al., 2014). However, some of these studies suffer from the limitation of using lipophilic dyes (e.g. PKH26, CMFDA) to label the MSCs (Jang et al., 2014; Xing et al., 2014), which have been reported to escape the labelled MSCs and can be internalised by the host cells, leading to false positive results (Li et al., 2013a).

In this study, we provide for the first time a direct and side-by-side comparison of the therapeutic potential and fate of UC-MSCs, BM-MSCs and A-MSCs in the refined mouse model of AKI induced by bilateral IRI. Using longitudinal transdermal renal clearance measurements and the serum biomarkers creatinine, blood urea nitrogen and cystatin-c as functional renal assessment, and histopathological scoring of both kidneys to determine kidney health, we analysed efficacy of the different MSC types. Furthermore, we monitored the presence and persistence of the three cell types using bioluminescence in vivo imaging longitudinally.

## 2. MATERIALS AND METHODS

### 2.1 Animals

Albino Black 6 (B6N-*Tyr^c-Brd^/*BrdCrCrl) male and female mice were purchased from Charles River, Italy, and used to generate an in-house colony maintained within the Biomedical Services Unit (BSU) at the University of Liverpool. Mice were housed in ventilated cages with a 12-hour light/dark cycle and access to water and food *ad libitum*. All animal experiments were performed in accordance with UK regulations set out in the Animals (Scientific Procedures) Act 1986 and following approval by the Animal Welfare and Ethical Review Board (AWERB) of the University of Liverpool and the Home Office, under project licences PPL70_8741 (‘Preclinical therapies for renal and cardiovascular injury’, approved 6.10.2015) and PP3076489 (‘Developing safe and efficacious cell-based therapies for kidney disease’, approved 2.11.2020). We report our work in line with the ARRIVE 2.0 guidelines (Percie du Sert et al., 2020). Animals were terminated using Schedule 1 methods, including rising CO_2_ levels or terminal anaesthesia using isoflurane. Humane endpoints throughout were weight loss of 30 percent or more during the acute injury phase, poor body condition, lack of appetite, unresolved skin inflammation, and necrosis in scarred tails after repeated tail vein injections

A total of 120 male B6 albino mice were used for the study presented here: 78 for experiments to assess MSC efficacy, 32 for experiments to assess MSC biodistribution, and 10 as uninjured and untreated control mice (**Supplementary Master Tables I, II, III**).

### 2.2 Surgical procedure

In 78 mice of the efficacy experiments, and 32 mice of the biodistribution experiments (**Supplementary Master Table I, II**), ischemia reperfusion injury (IRI) was induced in both kidneys by a dorsal approach as previously described (Harwood et al., 2022). Mice aged 9–10 weeks and weighing 20–27 grams were anesthetized with 2% isoflurane for 30 minutes and received 70 µL baytril, 500 µL saline, and buprenorphine (1mg/Kg) subcutaneously in the forelimbs prior to surgery.

Under continuous 2% isoflurane anaesthesia during the surgical procedure, mice were placed on a heat pad in prone position while the body temperature was monitored by a rectal probe and controlled by a homeothermic monitor system (PhysioSuite, Kent Scientific, Torrington) in the range of 36.5–37.0 °C. To exteriorize the kidneys, a small incision in the dorsal skin was made along the midline of the mouse. The kidneys were exposed and held with curved Taylor forceps while blunt forceps were used to release the fat tissue surrounding the renal pedicle. The renal pedicle was then clamped using a non-traumatic vascular clamp, with the clamp of the right kidney being set first followed immediately by clamp to the left kidney. Ischaemia was assessed by a change of colour in the kidneys from red to dark brown. During the clamping, the kidney was returned to the retroperitoneal space, covered with the muscle, and a sterile gauze soaked in sterile water was placed over the surgical incision to preserve moisture. Clamps were removed after 27.5 minutes, and reperfusion was visually confirmed by the kidneys immediately turning a red colour. The kidneys were returned to their space, and muscle and skin layers were repaired with absorbable 6-0 sutures.

After surgery, mice were transferred to a 37 °C warmed chamber for 30 minutes before returning to the housing cage. The appearance, weight and behaviour of mice were assessed daily to monitor welfare. Humane endpoints

### 2.3 Isolation of cells and culture

Umbilical cord (UC-) MSCs from two independent donors were obtained from the National Health Service Blood and Transplant (NHSBT, Liverpool, UK), licenced under the Human Tissue Act (HTA). The cells were isolated under good manufacturing practice. In brief, the cord was halved horizontally, cut into pieces of 2-3 cm in length and then cultured undisturbed for 7 days. The tissue pieces were subsequently removed and the adherent cells cultured for 2 passages. Bone marrow (BM-) MSCs, were provided by the University of Galway (Ireland), after being purchased from Lonza (Basel, Switzerland) and expanded for at least 2 passages. Adipose derived (A-) MSCs were provided by the University of Heidelberg (Germany), following the Mannheim Ethics Commission II approval (vote 2011-215N-MA)(Calcat et al., 2023). The cells were isolated from lipoaspirates harvested after informed consent. The tissue was digested with NB4 Collagenase (Serva/Nordmark), strained to remove the undigested tissue, and the cells were seeded and cultured for at least 2 passages. In all cases the cells were cryopreserved after the initial expansion and then shipped to the laboratory at the University of Liverpool for experiments. Once there, the cells were cultured following standard mammalian tissue culture protocols in MEM-α containing GlutaMAX (Gibco) supplemented with 10% foetal bovine serum (FBS; Gibco) at 37 °C in a humidified incubator, with 5% CO_2_. MSCs were routinely expanded and passaged when reaching 60-90% confluence.

### 2.4 Preparation of cells for in vivo administration

MSCs at passage 6 were used as a therapy. Cells were detached using trypsin/EDTA, centrifuged at 400g, and then 2.5×10^5^ MSCs were suspended in 100 µL sterile phosphate-buffered saline (PBS; Sigma). Cells were kept for up to 2 hours on ice until administration into animals. Prior to administration the cells were gently re-suspended and then injected with a 29G insulin syringe via the tail vein immediately after the surgical wounds were sutured.

For imaging experiments, MSCs were transduced with a lentiviral vector (LV) carrying a bicistronic construct encoding the luc2 firefly luciferase (FLuc) reporter and a green fluorescent protein, ZsGreen. The pHIV-Luc2-ZsGreen vector was a gift from Bryan Welm and Zena Werb (Addgene plasmid #39,196). Lentiviral particles were produced using standard protocols by co-transfection of HEK cells with the transfer vector (pHIV-Luc2-ZsGreen), an envelope plasmid (pMD2.G) and a packaging plasmid (psPAX2), concentration by ultracentrifugation and titration using HEK cells, based on ZsGreen expression (Amadeo et al., 2023a; Amadeo et al., 2022).

Transduced MSC populations were produced by infecting the cells overnight at 37 °C with a multiplicity of infection (MOI) of 5 in the presence of 6 μg/mL 40 kDa diethylaminoethyl-dextran (DEAE-dextran)(Amadeo et al., 2023b). Then, the medium was replaced, and the cells grown until 60-90% confluence before they were sorted based on ZsGreen fluorescence using a FACSaria II (BD Biosciences) to obtain the population of cells expressing the transgene (FLuc^+^ MSCs).

### 2.5 Experimental design of MSC efficacy studies

A total of 20 animals were assigned as cohort to each of three experiments assessing efficacy of UC-MSC Donor 1 vs PBS; BM-MSC vs PBS, or A-MSC vs PBS. In addition, 18 animals were assigned as cohort to the experiment assessing efficacy of UC-MSC Donor 2 vs PBS. Half of the animals in each cohort received 100 μL PBS (control group) and the other half received a single dose of either UC-MSCs Donor 1, UC-MSCs Donor 2, BM-MSCs, or A-MSCs by intravenous injection (iv) into the tail vein (cell therapy group). Based on previous experience in our lab regarding experimental feasibility (Harwood et al., 2022), experiments on an individual day were performed in batches of 8 (UC-MSCs Donor 2 batch 1) or 10 animals (UC-MSCs Donor 1, UC-MSCs Donor 2 batch 2, BM-MSCs, A-MSCs) consisting of equal PBS control and cell therapy-treated mice. IRI-injured mice were assigned following stratified randomisation by GFR to either PBS control (4 or 5 animals) or cell therapy group (4 or 5 animals), achieving similar GFR baseline values across groups (see 2.6.1). Administration of cells or PBS was performed in a blinded way, such that the operator was not aware of the substance administered.

The progression of AKI and injury severity were determined on day 1 and 3 post-IRI based on a transdermal device to longitudinally measure the GFR (see 2.6.1). Serum and kidneys were collected on day 3 after the intervention (**Supplementary Figure 1**). Animals were excluded from the study if: (a) any experimental conditions (including change of temperature during the surgery, for example by loss of the rectal temperature probe or temperature recording above 37.5 or below 36.5 °C, animal welfare, for example swollen feet) changed during the IRI procedure, (b) excessive bleeding occurred during the surgery, (c) abnormalities on the gross anatomy of kidney were observed before the procedure, (d) less than 50% of the therapy was administered intravenously, (e) severe injuries, trauma, or welfare concerns not related to the experiment were present. The number of animals excluded from the final analysis for each experiment and the rationale are reported. Therefore, the group numbers reported in the results are reflective of the applied exclusion criteria (see **Supplementary Master Table I**).

### 2.6 Measurements of kidney function and damage

#### 2.6.1 Glomerular filtration rate (GFR)

A transdermal mini GFR monitoring system (Medibeacon, Mannheim) relying on the clearance of FITC-sinistrin was used as previously described (Scarfe et al., 2018a). In short, mice were anesthetized (isoflurane 1.5%; 1.0 L/min, O_2_) for the device placement, background acquisition and intravenous (IV) injection of FITC-sinistrin. The device was placed on the pre-shaved skin of the mouse, secured with silk tape and left for 5 minutes to measure the background signal. Next, 0.075 mg/g body weight of FITC-sinistrin (20 mg/mL in PBS) (Medibeacon, Mannheim) was injected via the tail vein. Mice were allowed to recover from the anaesthesia and left in single cages with access to food during the 90 minutes of measurement after which the device was removed from the animals. For all 78 animals of the IRI cell therapy studies, baseline readings were obtained around a week before the IRI surgery.

The data were analysed with MB Studio software version 2.5, where the half-life of FITC-Sinistrin (t_1/2_) was obtained by using a three-compartment linear model and transformed into GFR data by using a semi-empirical conversion factor corrected for body weight (bw) as described by Schreiber and colleagues (Schreiber et al., 2012) (Equation 1).

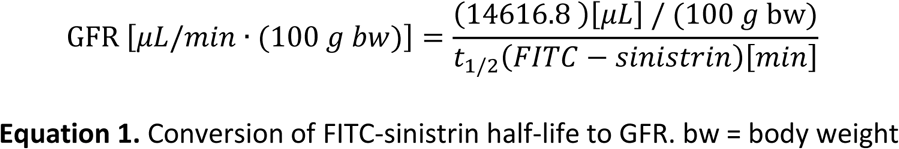

To classify kidney injury based on the RIFLE criteria, we determined the percentage change in kidney function by calculating the percentage change compared to baseline GFR for each individual animal (Bellomo et al., 2004; Harwood et al., 2022).

#### 2.6.2 Serum biomarkers

On day 3 post IRI and after the final transdermal GFR assessment, blood was collected under terminal anaesthesia via cardiac puncture with a 25G needle introduced from underneath the diaphragm. Upon blood collection, animals were individually euthanized by cervical dislocation and the samples were allowed to clot at room temperature for 1-2 hours. Blood samples were then centrifuged at 2000g for 20 minutes, and the supernatant was carefully collected and centrifuged at 75g for 4 minutes. Serum was aliquoted and stored at −20 °C until use. Sera from 10 uninjured healthy male Bl/6 albino mice were collected and used as normal reference values and as internal controls for the assays (see **Supplementary Master Table III)**.

Freeze-thaw cycles were avoided and serum was processed and analysed according to the manufacturer’s guidelines of the marker assays: Serum creatinine was determined by Jaffe’s reaction (Detect X Serum Creatinine Detection Kit, Arbor Assays, Ann Arbor, MI, USA); blood urea nitrogen (BUN) was measured by the Jung colorimetric method (QuantiChrom Urea Assay Kit, BioAssay Systems, Hayward, CA, USA); and cystatin-C was assessed by solid phase sandwich ELISA (Quantikine ELISA, R&D Systems, UK). From the 10 uninjured mice we could not obtain serum data for all three biomarkers (see **Supplementary Master Table III)**.

#### 2.6.3 Histology

Mouse kidneys from animals which had been taken through to the end of the experiment, were fixed in 10% neutral buffered formalin for 24 – 72 hours. After fixation both kidneys were sectioned sagittally in two halves with a kidney slicer manufactured in-house. The specimens were then immediately returned to formalin until processing. The specimens were paraffin-embedded and 3 μm sections were collected from left and right kidneys, and stained with Periodic Acid-Schiff (PAS) for histological analysis.

Histological examination was performed blindly by an experienced board-certified veterinary pathologist (LR), using an adapted semi-quantitative scoring system described by Wang and colleagues (Sharkey et al., 2019; Wang et al., 2005). For each animal, left and right kidneys were assessed by scoring one section each from both halves of left and right kidneys. Histological damage was scored in the cortex and outer stripe of outer medulla (OSOM) by counting the percentage of four typical histopathological characteristics of IRI that included: 1) cell necrosis, 2) loss of brush border, 3) cast formation, and 4) tubule dilation (**Supplementary Figure 2**). A total of 10 fields of view (200x) were randomly examined for each slide and scored for damage as follows: **0** = no damage, **1** = 1-25%, **2** = 26–50%, **3** = 51–75%, **4** = 76–100% damage. Mean Wang scores were calculated from left and right kidneys separately, and by combining left and right from each animal.

### 2.7 Biodistribution of FLuc^+^ MSCs in mice with IRI

MSCs expressing firefly luciferase (FLuc^+^ MSCs) were tracked by bioluminescence imaging (BLI) in healthy and injured mice. Because combining IRI surgery, cell administration and BLI on the same day may compromise animal wellbeing, three independent experimental groups were used. In this independent experiment, group 1 comprised mice that had successfully undergone IRI, were injected with the respective cells on day 0 (UC-MSC Donor 1: 4 animals; BM-MSC: 3 animals; A-MSC: 3 animals) and subsequently imaged by BLI under terminal anaesthesia immediately after surgery (**Supplementary Figure 3, Supplementary Master Table II**). In group 2, mice that had undergone IRI surgery and were injected with the respective cells (UC-MSC Donor 1: 5 animals; BM-MSC: 4 animals; A-MSC: 4 animals), were allowed to recover and imaged at days 1, 3 and 7. GFR was measured in group 2 animals before the surgery/cell administration and then on days 1 and 3. Group 3 which comprised mice (n = 3 animals per MSC type) that had not undergone the IRI procedure but received the cells on day 0, was included as a no-injury control (**Supplementary Figure 3**). To account for any possible light attenuation during the imaging due to the presence of the fur we shaved the back of the IRI and control animals.

At the end of the surgical procedure and in the same anaesthesia session, 2.5×10^5^ of the following cell types were injected in 100 μL of sterile saline in the tail vein: FLuc^+^ UC-, FLuc^+^ BM-, or FLuc^+^ A-MSCs. Control (uninjured) animals were anaesthetised with 2% isoflurane and received the cells in the same way. For the acquisition of BLI data, mice received a subcutaneous administration of 200 μL of 47 mM D-Luciferin (Amadeo et al., 2022) and, 20 minutes later, were imaged using an IVIS Spectrum system (Perkin Elmer) in dorsal position. The acquired signal was normalised to radiance (photons/second/centimeter^2^/steradian, p/s/cm^2^/sr) and the signal originating from the thoracic area of the animals was quantified using the region of interest (ROI) tool of the IVIS software (Living Image v. 4.5.2) to obtain the total number of photons emitted in that specific area and displayed as total flux (photons/s). Each imaging session was performed using the following settings: open filter, binning of 8, f-stop of 1, and 60 seconds exposure time on day 0, and 180 seconds exposure time on days 1, 3 and 7.

### 2.8 Statistical analysis

Differences in GFR measurements over time between MSC-treated and PBS-treated mice were assessed using linear mixed effects models. Since an initial drop in GFR was followed by a rise, a piecewise model was considered with separate time slopes for the period between baseline and day 1, and the period between day 1 and day 3. To investigate whether change in GFR levels over time was different between animals who received PBS treatment and those receiving MSC-treatment, we included an interaction term between time and treatment. We considered all GFR measurements within a single model and included a factor in the model to account for differences in PBS groups across experiments (A-MSC vs PBS, UC-MSC (donor 1) vs PBS, UC-MSC (donor 2) vs PBC or BM-MSC vs PBS).

To discern statistical differences in data between PBS-treated and MSC-treated IRI mice, based on the analysis of serum markers and histology on day 3 after IRI, we considered linear regression models, with factors for treatment and experiment type. BUN values were log transformed to ensure normality of model residuals. To aid interpretation, independent t-tests and paired t-tests were applied to compare histological Wang scores between PBS-treated and MSC-treated groups, without adjustment for multiple comparisons.

All statistical analysis was performed using GraphPad Prism version 10 (GraphPad Software, California-USA) and R version 3.4.2. A p-value (p) of less than 0.05 (*p < 0.05) was considered to be statistically significant, otherwise it was declared nonsignificant.

## 3. RESULTS

In order to determine whether MSCs obtained from human umbilical cord, bone marrow or adipose tissues had different efficacy in protecting or ameliorating AKI, we performed three assessments for each type of MSC in the same experiment, using a well-established bilateral kidney IRI mouse model of AKI. Kidney function was assessed by transdermal measurement of GFR (Scarfe et al., 2018a; Schreiber et al., 2012), and the urine biomarkers serum creatinine, blood urea nitrogen (BUN) and cystatin-C (Harwood et al., 2022), while kidney health was determined using histopathological scoring. Not all mice contributed to each analysis. Reasons for exclusion are listed in **Supplementary Master Table 1**.

### 3.1 Effect of MSC-based therapies on kidney function as assessed by GFR

For a longitudinal assessment of kidney function, GFR measurements were collected from the same animal on the day before the IRI surgery (baseline = day 0), and on days 1 and 3 after the surgery. We included UC-MSCs from two independent donors in this analysis to robustly confirm results.

GFR measurements revealed longitudinal dynamic changes in kidney function for each animal group (**Figure 1A-D**, **Supplementary Table 1, 2**). We observed that injured animals treated with BM-MSCs or A-MSCs had a 25% higher mean GFR on day 3 compared to respective PBS-treated mice. By contrast, injured animals treated with UC-MSCs from donor 2 had a reduced mean GFR compared to the PBS-treated group on day 3, while injured animals treated with UC-MSCs from donor 1 had a similar mean GFR on day 3 to the PBS-treated group. It is important to note that the group sizes were relatively small and variability quite large, leading to large standard errors and confidence intervals.

**Figure 1.**
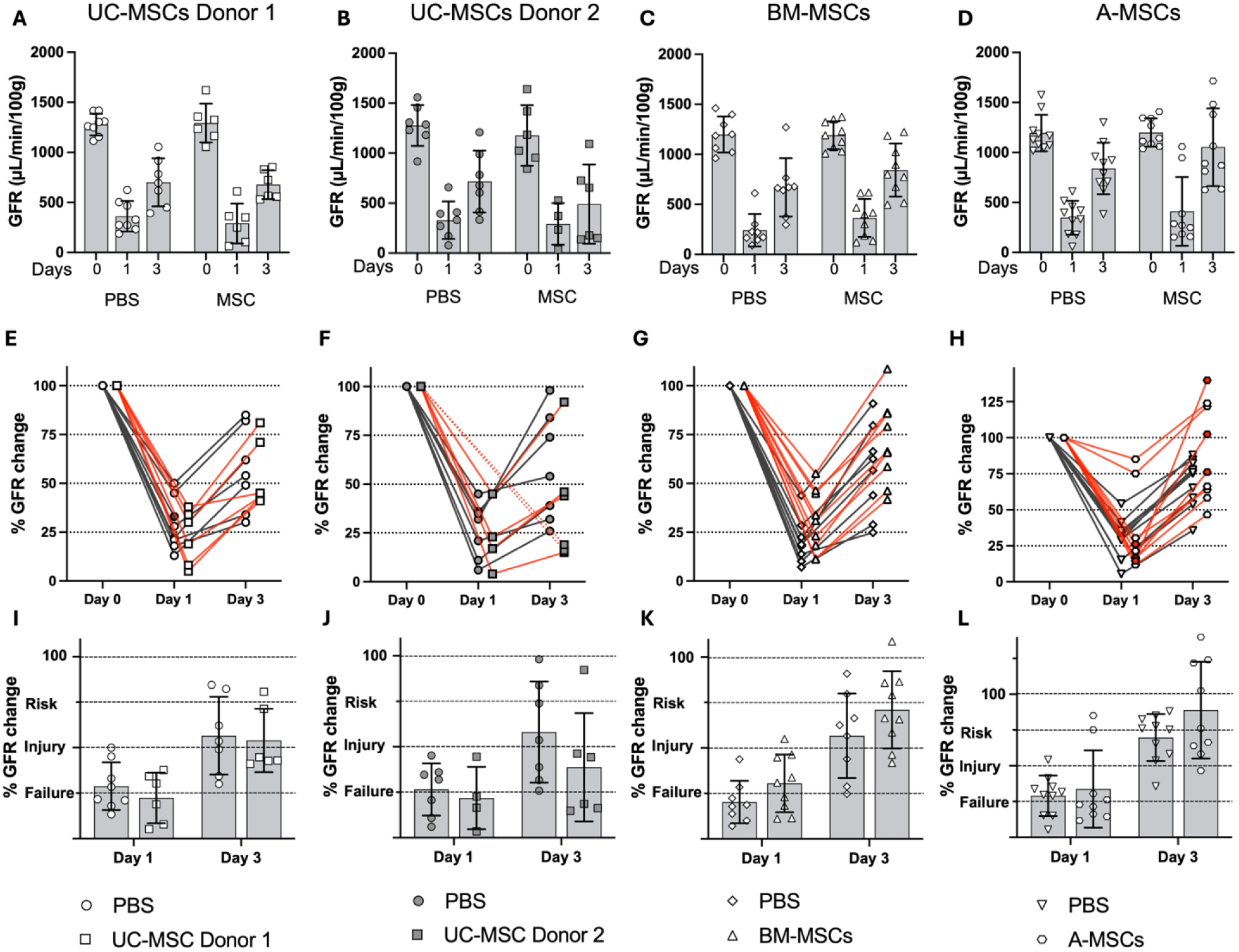
Longitudinal changes in kidney function before and after renal IRI and in response to MSC treatment. **(A-D)** Transdermal GFR indicated that in most animals, baseline values (day 0) were above 1100 µL/min/100g which decreased by day 1 and had partially recovered by day 3 post IRI. After treatment with either UC-MSCs (A, donor 1, n=6, or B, donor 2, n=6), BM-MSCs (C, n=9) and A-MSCs (D, n=9), GFR values were compared with those in control groups (n=7-10). **(E-H)** Individual mouse data over time according to percentage of kidney function (% GFR change compared to day 0). Grey data point in E indicates animal where no recording could be made at day 3. Red data points in H indicate kidney function in or near failure at day 1 and recovered at day 3. **(I-L)** Based on the percentage of kidney function, individual mice were classified using the RIFLE criteria. At day 1, most animals were in the injured or failure category. By day 3, most animals had recovered between 10-45% of their kidney function. Data are displayed with mean ± SD in (A-D, I-L).

We applied the RIFLE criteria (Bellomo et al., 2004; Harwood et al., 2022) to classify the grade of severity in AKI, thus providing clinically relevant context for the mouse GFR data. On day 1, kidney function based on GFR had decreased to around 20-30% of the baseline value in all conditions, independent of the MSC or PBS treatment (**Figure 1E-L**, **Supplementary Table 2**). This led to animals reaching risk, injury or failure categories (**Figure 1I-L**). Linear mixed models revealed that for animals in the PBS control groups the observed decrease in GFR values between baseline and day 1 was significant, followed by a significant increase between day 1 and day 3 (**Supplementary Table 3**). No significant difference was observed in baseline levels between cell-treated animals and PBS controls.

Whilst each group experienced a drop in GFR followed by a subsequent rise, we found no evidence of different trajectories between the different treatment groups.

Additional observations of the effects of A-MSC administration included that 44% (4/9) of mice that had received A-MSCs did not go into kidney injury or failure and showed normal kidney function on day 3, suggesting a protective effect of the A-MSCs for these mice. Furthermore, 33% (3/9) of mice that had received A-MSCs recovered normal kidney function (75% or above) on day 3 after being in or close to failure on day 1 (**Figure 1H**). These degrees of improvements in kidney function were not seen in any of the control groups or animal cohorts treated with UC-MSCs or BM-MSCs. Because we failed to see any statistically significant difference in efficacy in the UC-MSCs from the two different donors, we focussed on the UC-MSCs from donor 1 from here onwards.

In all cases, the acute kidney injury induced by IRI led to body weight loss that the mice had not recovered from by the end of the experiment (**Supplementary Figure 4**). Animals after IRI and A-MSC treatment appeared to be better protected against weight loss at day 1, although at day 3, these animals had reached similar weight loss to the PBS controls in this cohort.

### 3.2 Effect of MSC-based therapies on kidney function as assessed by serum biomarkers

As a second measure of kidney injury, we assessed the serum biomarkers creatinine, BUN and cystatin-C in blood samples collected at the end of the experiment (day 3). Mean creatinine levels in the BM-MSC group were significantly lower (by 21.8%) than in the cohort controls, and similar to, but slightly lower than, the mean levels of uninjured reference animals (**Figure 2A**, **Supplementary Table 4, Supplementary Master table III**). By contrast, mean creatinine levels in the UC-MSC group were 13.3% lower than cohort controls but without statistical significance, and in the A-MSC group similar to the cohort controls. Linear regressing models confirmed these results (**Supplementary Table 5**). Currently, it is not clear whether this observation has any biological significance.

**Figure 2.**
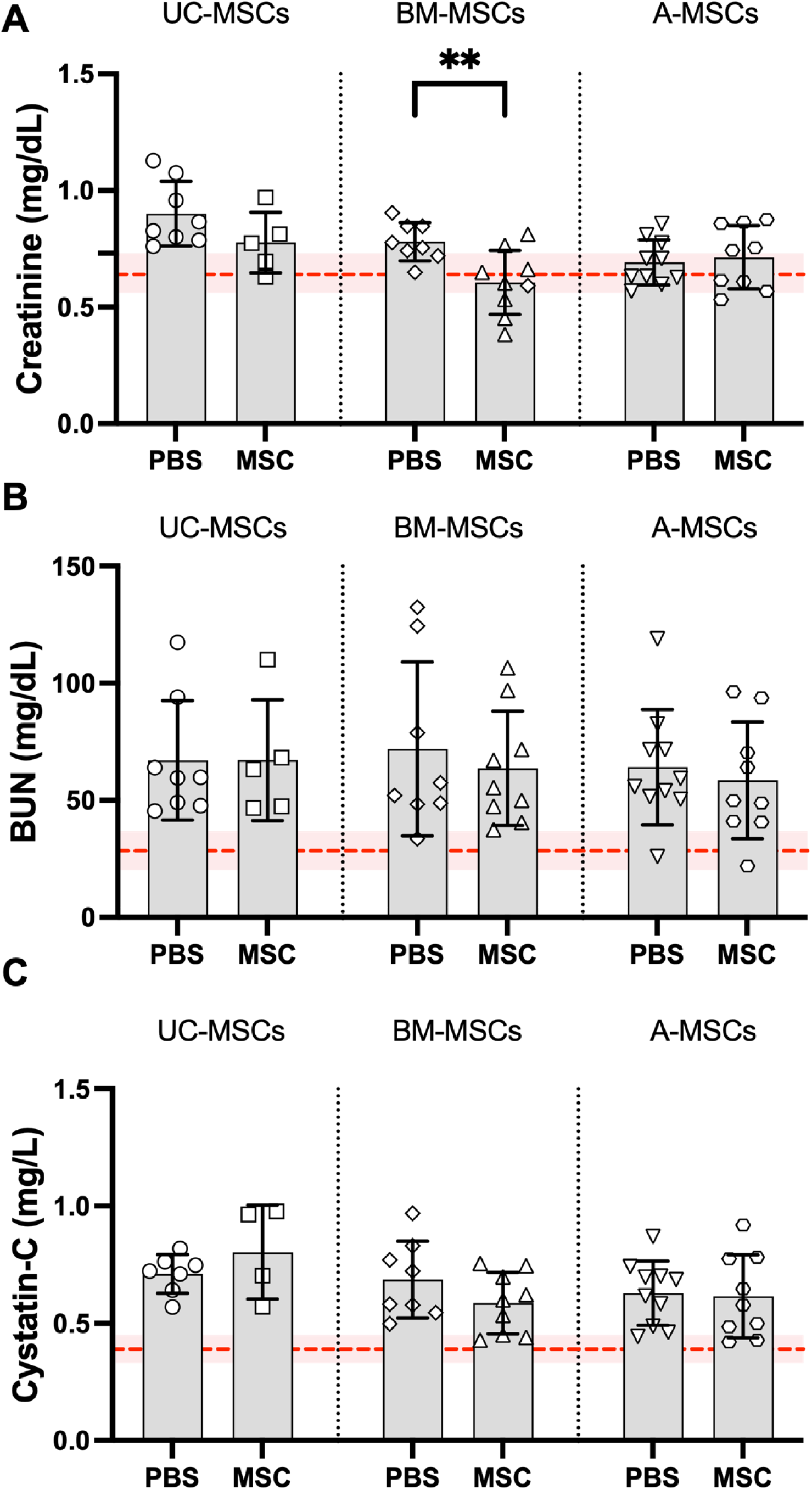
Levels of serum markers in mice 3 days after IRI and MSC treatment. **(A)** Serum creatinine, **(B)** BUN and **(C)** Cystatin-C were measured in animals on day 3 after IRI, presented by treatment cohort. **(A)** Mean serum creatinine level was lower in the UC-MSC treated animals (n=5) compared to PBS controls (n=8), and in the BM-MSC-treated animals (n=9) compared to cohort controls (n=8). For BM-MSC treatment, this difference was statistically significant (p=0.005). There was no difference in the means for the A-MSC-cohort between cell-treated (n=9) and PBS control mice (n=10). **(B)** Mean BUN levels were slightly lower in all groups with MSC treatment (UC-MSC: n=5; BM-MSC: n=9; A-MSC: n=9) compared to PBS controls (UC-MSC: n=8; BM-MSC: n=8; A-MSC: n=10). **(C)** Mean cystatin-C level was lower in the BM-MSC treated animals (n=9) compared to PBS controls (n=8), but this was not statistically significant. There was no difference in the means for the UC-MSC- and A-MSC-cohorts between cell-treated (UC-MSC: n=4; A-MSC: n=9) and PBS control mice (UC-MSC: n=7; A-MSC: n=10). Bar chart depicting individual animal values with mean and SD, per treatment group within cohorts. For comparison purposes, the means of measurements from serum from healthy, uninjured animals (n=10 for Serum Creatinine; n=6 for BUN; n=7 for Cystatin-C) and 95% CI are displayed as red stippled line and pink area, respectively.

We detected slightly lower means of BUN values in the mice of BM- and A-MSC-treated groups when compared to their respective PBS-treated controls (**Figure 2B**). However, no statistically significant differences were found (**Supplementary Tables 4, 5**). Values from all injured animals with or without cell-treatment were higher (means between 58 and 72 mg/dL) than the mean of the uninjured reference group (28 mg/dL) (**Supplementary Table 4**).

Mice in the groups that had received UC-MSCs or A-MSCs had cystatin-C levels higher or comparable to their PBS-treated controls, respectively (**Figure 2C**). By contrast, animals in the BM-MSC group had slightly lower values compared to the PBS-treated control group. Differences in cystatin-C means between the cell-treated groups and controls were not statistically significant for any of the experiments (**Supplementary Tables 4, 5**). Values from all injured animals were higher (means between 0.59 and 0.75 mg/L) than the mean of the uninjured reference group (0.39 mg/L) (**Supplementary Table 5**). All data that have contributed to this analysis are provided in **Supplementary Master Tables I and III.**

### 3.3 Histopathological analysis and comparison of the beneficial effects of MSCs in mice with IRI

To detect the presence of cell necrosis, loss of brush border, cast formation, and tubular dilation in tissues collected from the mice at day 3, we performed PAS staining of kidney sections from one (BM-MSC and A-MSC) or both batches (UC-MSC) of animals per MSC-treatment, and scored left and right kidney sections (**Supplementary Master Table I**). Representative images of reference and injured renal specimens demonstrate differences in renal tissue integrity in kidneys from both PBS-and MSC-treated mice (**Supplementary Figure 5)**.

We observed that in two animals after treatment with UC-MSCs, the right kidneys showed prominent evidence of diffuse coagulating necrosis and the presence of intravascular fibrin thrombi, which led us to exclude these from the histological analysis (**Supplementary Master Table I**). These morphological changes were not expected in an IRI model, but could be due to thrombosis as a consequence of the experimental approach. In one of the two animals, we had observed a lack of perfusion of the right kidney at the time of skin wound closure (see comment in **Supplementary Master Table I**), which might have a causal relationship to the histopathology in this kidney.

The combined mean Wang scores for left and right kidney sections of the UC-MSC- and BM-MSC-treated animals were slightly lower than those of the respective PBS-treated control animals (**Figure 3A, Supplementary Table 6)**. In contrast, based on the mean combined Wang score, mice that had received A-MSCs presented slightly more histological damage than those of the control group. Overall, there were no statistically significant differences between any corresponding cell therapy and control groups when scores for right and left kidneys were combined for each animal (**Supplementary Table 7**).

**Figure 3.**
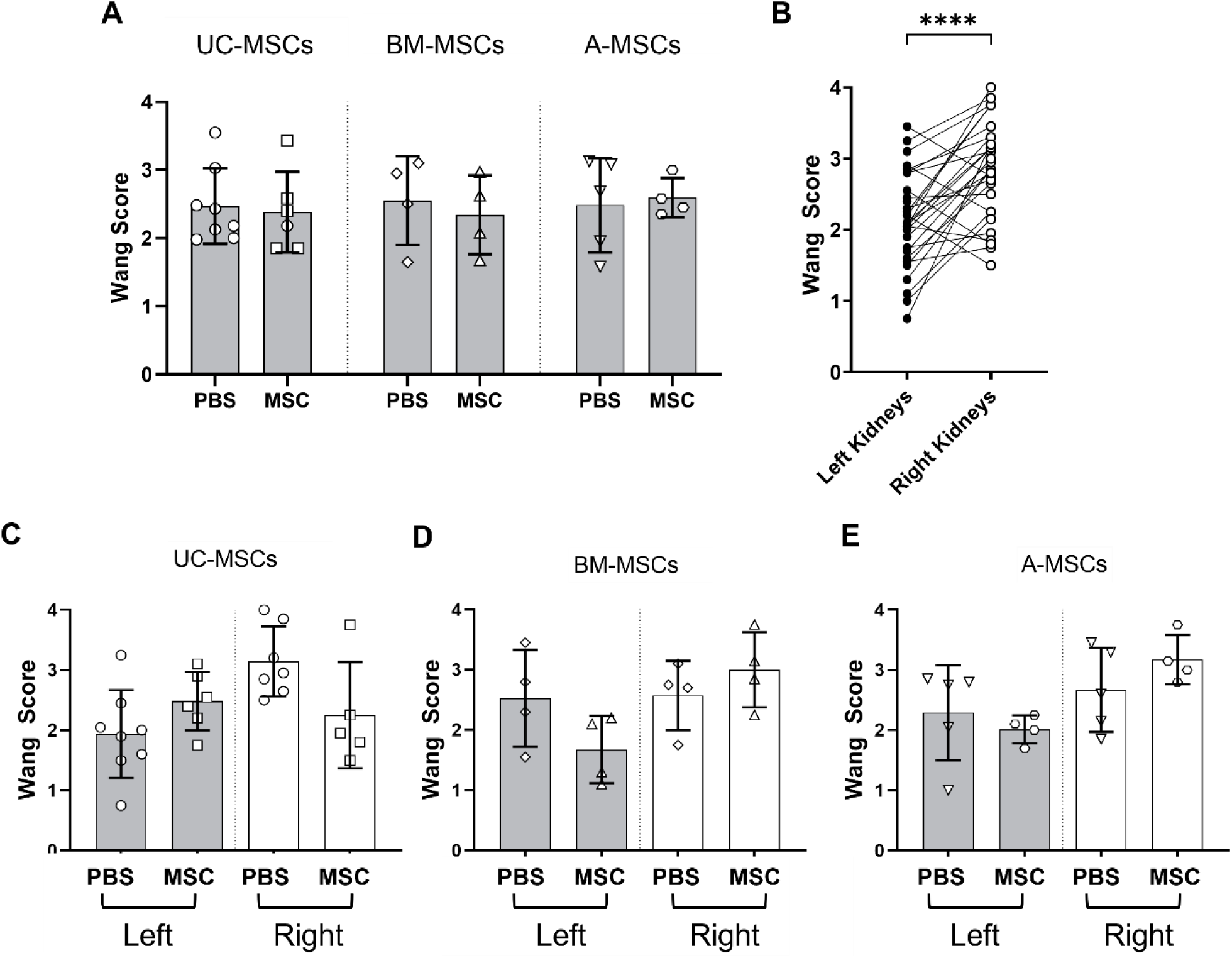
Histological analysis using the Wang modified system of kidney sections stained with Periodic acid-Schiff (PAS) 3 days after IRI and MSC treatment. **(a)** Combined mean Wang scores for left and right kidneys for each animal analysed after IRI and administration of UC-MSCs (left, n=5), BM-MSCs (centre, n=4) or A-MSCs (right, n=4) or respective control groups (n=4-5). Note that one of the data points for UC-MSC-treated mice consists of only the score for the left kidney since the right kidney displayed diffuse coagulative necrosis (see SI Master Table I). **(b)** Mean Wang scores for all left (n=31) and all right (n=29) kidneys independent of treatment after IRI indicated that right kidneys were significantly more injured. **(c-e)** Wang scores for left and right kidneys separately of UC-MSC-(c), of BM-MSC- (d) or A-MSC- (e) treated animals compared with PBS-treated animals. Data are displayed with mean ± SD.

When we analysed the Wang scores for left and right kidneys separately, we observed that the mean Wang score of the left kidneys of all animals, irrespective of cell therapy treatment or not, was significantly lower than that of the right kidneys (**Figure 3B**). This was likely due to the order in which the surgical clamping was performed in the mice. Subsequently, we plotted the Wang scores of sections from left and right kidneys separately for all animals. This revealed differences between left and right kidneys within treatment cohorts by cell type, but without any significance when compared to the injured control animals (**Figure 3C-E**). These results suggest that any possible biological relevance of A-MSCs or BM-MSCs in ameliorating kidney function after kidney IRI, as detected by GFR or BUN measurements, was not reflected in the kidney histopathology of these mice. It is important to note that the low sample sizes in all groups were a limitation to drawing conclusions regarding biological relevance and efficacy of the MSCs used.

In summary, based on the three different measures of kidney function and health, transdermal GFR, serum biomarkers and histology, our data indicate that the three different MSC types have modest if any effects in ameliorating kidney injury induced by the bilateral IRI model of AKI. The only significant functional effect observed, was based on serum creatinine in animals administered BM-MSCs.

### 3.4 Tracking the fate of the MSCs in mice after IRI

Next, we investigated whether the presence of kidney injury had an impact on the migration or survival of UC-, BM- and A-MSCs. For this purpose, we used BL imaging to track the biodistribution and persistence of the administered cells in animals after IRI, building on previously established protocols (Amadeo et al., 2023a)(see also **Supplementary Master Table II**). The experimental design incorporated three different groups of animals: Animals in group 1 underwent IRI surgery, MSC administration and immediate BLI under terminal anaesthesia, while animals in group 2 were allowed to recover after IRI and MSC administration, followed by BLI on days 1, 3 and 7. Animals in group 3 were not injured but still received MSCs followed by BLI on days 1, 3 and 7 (**Supplementary Figure 5**). We monitored kidney function in animals from group 2 using longitudinal GFR measurements on days 0, 1 and 3, and observed similar dynamics in the functional injury response to those reported above (**Figure 4A**, see **Figure 1**): on day 1, GFR values had dropped and on day 3, recovered to varying levels, but were consistently below the baseline values.

**Figure 4.**
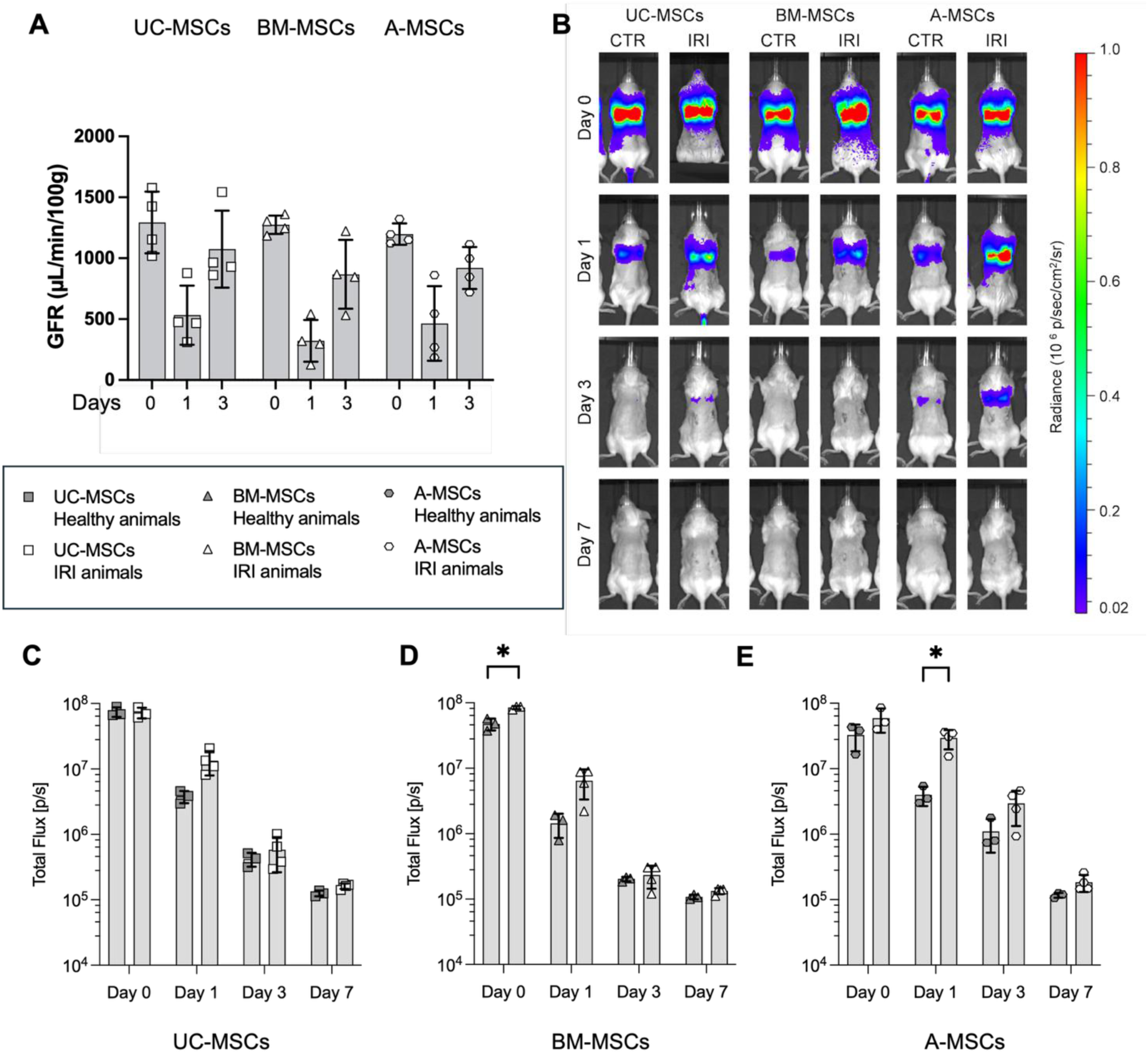
MSCs were sequestered in the lungs of healthy and IRI animals where they died. **(A)** Longitudinal GFR measures confirming the injury response in animals after IRI and with MSC treatment (n=4 for all MSC-treated cohorts). Baseline transdermal GFR values (day 0) decreased by day 1 and had partially recovered by day 3 post IRI. The response in all groups was similar to that shown in Figure 1 A-D. **(B)** Representative bioluminescence images of control and IRI mice up to 7 days post administration of FLuc^+^ UC-, BM- and A-MSCs (radiance scale from 0.2×10^5^ to 1×10^6^ p/s/cm^2^/sr). **(C-E)** Light output (p/s) as a function of time for each MSC type, incorporating three groups of mice: group 1 mice (n=3 for each MSC type) underwent IRI, MSC administration and immediate BLI under terminal anaesthesia, group 2 mice (n=4 for each MSC type) underwent IRI and MSC administration but were imaged on days 1, 3 and 7; group 3 mice (control, n=3 for each MSC type) were only administered MSCs and imaged on days 0, 1, 3 and 7. Significant differences were found for signals on day 0 in BM-MSC-injected cohorts between injured and non-injured mice (p=0.03, Brown-Forsythe and Welch ANOVA test, multiple comparisons), and on day 1 between injured and non- injured mice in the A-MSC-injected cohort (UC-MSCs: p=0.04, Welch’s t-test; A-MSCs: p=0.04, Brown-Forsythe and Welch ANOVA test). Graphical display as mean ± SD.

The bioluminescence signals obtained from healthy (CTR) and IRI animals (group 1) on the day of MSC administration indicated that immediately after injection, the cells were sequestered within the lungs (**Figure 4B**). In animals (groups 2 and 3) that were imaged on day 1 after cell administration, the bioluminescence signals were weaker in both the injured animals and the healthy animals compared to signals obtained on the day of administration (**Supplementary Master Table II**, **Supplementary Table 9**). At this stage the qualitative signal analysis in all animals indicated that cells remained confined to the lungs, independent of injury (**Figure 4B**). Interestingly, comparison of qualitative signal distribution and quantitative signal intensity (mean total flux) between healthy and IRI animals with the same cell treatment revealed that the signals in mice after IRI appeared to be stronger than in healthy animals, indicating that more cells may have persisted in the lungs of injured mice (**Figure 4C-E**). Mean total flux at day 1 was statistically different between control and IRI-injured mice after A-MSC injection (unadjusted for multiple comparison; p=0.04). Furthermore, 3 days post cell administration, signals from A-MSCs were still clearly detectable in the lungs, while only weak signals were detectable in some of the mice that had received UC-MSCs (**Figure 4B, E**). Finally, on day 7 no signals were detectable in any of the mice that had received cells (**Figure 4B-E**). Because signals were indistinguishable from background signal levels, this indicates that all injected cells had died. It is important to note that bioluminescence signals were only detected in the lungs and in no other organ, in animals of either of the groups in any of the imaging days, irrespective of the type of administered MSC.

## 5. DISCUSSION

Here, we have presented for the first time a direct side-by-side comparison to test the efficacy of human MSCs from the umbilical cord, bone marrow and adipose tissue, in ameliorating AKI in a bilateral kidney IRI mouse model. To quantify kidney function and health we used longitudinal transdermal GFR assessment, serum biomarkers, and histology. Based on our data neither MSC type demonstrated comprehensive efficacy by ameliorating kidney function and health across all measures.

Although we observed that on day 3 the means of transdermal GFR measurements in mice with IRI and administered with BM- and A-MSCs were 25% higher compared to the injured control animal groups, these results were not statistically significant. Furthermore, BM-MSC administration resulted in the lowest serum creatinine value on day 3, which was statistically significant compared to that of injured control mice. Histological scoring of the kidney tissues revealed that none of the MSC types caused a meaningful and significant reduction in the histological Wang score the kidneys compared to injured control mice. Overall Wang scores for all right kidneys were higher, indicating that they were more severely damaged than the left. It is important to mention that weights of animals with or without MSC therapy failed to recover in the 3-day experiments, suggesting that the overall wellbeing of the mice was affected by the bilateral IRI which could not be ameliorated by the administered cells.

Lastly, we compared the persistence and distribution of the three MSC types in mice with IRI by bioluminescence imaging and found that the cells persisted on day 1 significantly more in injured animals administered with UC-MSCs and A-MSCs when compared to mice without injury, suggesting a delay in clearing the cells. In all cases, the MSCs were sequestered in the lungs and failed to home to the kidneys.

While preclinical studies have revealed some levels of efficacy in a range of kidney injury models using different MSC types (Cai et al., 2014; Cao et al., 2010; da Silva et al., 2015; Fang et al., 2012; Geng et al., 2014; Hauser et al., 2010; Jang et al., 2014; Morigi et al., 2008; Shih et al., 2013; Wang et al., 2024a; Wang et al., 2020), a major breakthrough in clinical application is still outstanding (Amadeo et al., 2021; Fazekas and Griffin, 2020; Wang et al., 2024b). Understanding the mechanisms by which MSCs as cellular therapies confer protection, and promote repair and regeneration is a critical aspect of developing efficacious therapeutic interventions. Several mechanisms of action have been proposed for the MSCs to exert their therapeutic functions, such as direct and indirect immunomodulation (Zhou et al., 2019), including via the secretion of paracrine factors (Merimi et al., 2021) and extracellular vesicles (Almeria et al., 2022; Skovronova et al., 2021) which can carry microRNAs (Eleuteri and Fierabracci, 2019; Phinney et al., 2015). There is an ongoing debate whether MSCs need to be alive to exert their immunomodulatory effects, as apoptosis has been proposed as a mechanism by which the MSCs can interact and prime the host immune system (de Witte et al., 2018; Galleu et al., 2017; Gavin et al., 2019; Shokoohmand et al., 2025).

Our experimental approach to induce kidney injury by bilateral clamping of the kidneys was effective, since GFR in the mice was reduced by at least 50% on day 3 post injury, while serum markers were raised and considerable histological damage was detectable, as in our previously established model (Harwood et al., 2022). Since we used a model of acute injury, it was expected that the tissue damage resolved with time, which is seen by the partial recovery of kidney function within the time frame of the experiments (three days). Indeed, we observed that 27.5 minutes of bilateral clamping produced a reversible AKI since the transdermal GFR declined on day 1 but partly recovered by day 3.

The GFR data demonstrated that in our injury model, a clear therapeutic effect by any of the MSCs used was not detectable. We assessed GFR longitudinally which allowed us to track kidney function without the need for multiple blood sampling in the same animal over time. Furthermore, we stratified the mice using the RIFLE criteria according to their kidney function based on the transdermal GFR data. This experimental approach demonstrated no overtly significant improvement of the kidney function in animals that received MSCs after IRI, on day 1 nor day 3, which we confirmed using a linear mixed model analysis, however, group sizes were quite small.

Assessment of serum biomarkers revealed that BM-MSC administration led to significantly, ∼22% lower serum creatinine values at day 3 when compared to the injured control animals, which could indicate an improvement in renal function. However, we failed to detect any statistically significant improvement in BUN and cystatin-C, suggesting that the reduced creatinine levels observed for the BM-MSC treated group were likely to be of limited biological relevance. Two studies using bilateral IRI in mice have assessed the efficacy of human UC-MSCs, but either administered the cells 24 hours before (Jang et al., 2014) or after the surgery (Li et al., 2013b); in these studies, serum creatinine values were reduced by 1/3 to 1/2 on days 2 or 3 post injury. Similar results were obtained in a bilateral IRI mouse study, where the administration of BM-MSCs 24h post-surgery led to a ∼50% reduction in serum creatinine values on day 2 post injury (Xing et al., 2014). There are clearly differences in the experimental design of these studies to the one presented here, since we administered the cells immediately after surgery. A study by Wise and colleagues administered human BM-MSCs in a mouse model of bilateral IRI immediately after injury, similar to our approach, and detected a reduction of serum creatinine by 1/3 on day 3 post injury (Wise et al., 2014). Thus, a number of studies have demonstrated reno-protective effects of human BM- or UC-MSCs in murine bilateral IRI models using serum biomarkers, while we did not find convincing evidence of efficacy. Although mice that were administered human BM-MSCs (but not A-MSCs or UC-MSCs) displayed a significant reduction in creatinine, the lack of improvement in the other parameters tested suggests that the reduction in creatinine has little biological relevance. Currently, the discrepancy between our results and those of the aforementioned earlier studies is not clear. However, we used three different measures including longitudinal GFR measurements to assess kidney health and function, yielding consistent findings. Longitudinal GFR measurements have not been reported in other studies that assessed the efficacy of cells as therapies. This raises the question of whether the proportion of positive findings reported in the literature may be a consequence of publication bias.

In our experimental design, we administered a single dose of 2.5×10^5^ MSCs immediately after the surgical induction of the injury. We believe that the cell dose we applied is an adequately moderate dose for this experimental model, particularly due to our experience that higher cells doses can lead to adverse effects e.g. potential intravascular cell aggregation/clotting and embolism (unpublished observation). Previous studies administered 1×10^6^ cells and reported efficacy (Jang et al., 2014; Li et al., 2013b; Wise et al., 2014; Xing et al., 2014). However, efficacy has also been observed in a mouse IRI model following administration of 1×10^5^ human MSCs, as evidenced by significantly reduced serum creatinine and BUN levels at day 2 after surgery (Ranghino et al., 2017). Therefore, our dose was within the range reported in previous studies.

Histological analysis of the kidneys in our study demonstrated a slight mean reduction in the Wang score when mice had been treated with UC-MSCs or BM-MSCs compared to injured control mice. However, this difference was not statistically significant when left and right kidneys per treatment were pooled. Interestingly, we observed that on average the right kidneys presented a significantly higher degree of tissue damage after IRI than the left kidneys. The physiological basis for this observation is not clear although the order of setting the clamp (first right, followed immediately by left) could be a contributing factor to the generally higher tissue damage in the right kidneys. Importantly, animals treated with UC-MSCs had a not-significantly reduced injury in the right kidney compared to the right kidneys of control animals, suggesting that these cells could ameliorate the histological tissue damage. Although previous studies have demonstrated that cell therapies can reduce tissue damage in kidney IRI models, none of these studies have indicated if there were consistent differences in the degree of damage between left and right kidneys (Jang et al., 2014; Li et al., 2013b; Wise et al., 2014; Xing et al., 2014).

Previous studies have reported efficacy of systemically administered MSC therapies in rodent kidney models, irrespective of whether the cells populate the kidneys or are entrapped in the lung vasculature (Li et al., 2013b; Rota et al., 2018; Schubert et al., 2018; Shih et al., 2013). In the current study, bioluminescence analysis of the biodistribution of the three MSC cell types in animals after IRI showed that the cells sequestered in the lungs irrespective of the MSC type and the presence of the kidney injury. The bioluminescence images revealed that the MSCs failed to migrate to the injured kidneys, neither on the day of administration nor on any of the following days. The animals with IRI showed a similar GFR response to administration of the three cell therapies as the animal cohorts in Figure 1. We also observed that the MSCs died soon after administration since on day 1 around 90% of the signal in the healthy control animals had been lost. Interestingly, mice with an injury had slightly higher bioluminescence signals on day 1 than the corresponding control healthy control animals, irrespective of the type of MSCs. This observation suggests that the MSCs persisted in the lungs of injured mice slightly longer than in healthy animals. The reasons for this difference in cell persistence are unclear but could be linked to the inflammatory state induced in the injured animals (Amadeo et al., 2023a; Luk et al., 2016; Pichardo et al., 2022); of note, the animal group sizes were very small. Nonetheless, seven days after cell administration no signal was detected in any of the animals, confirming that the cells disappeared regardless of kidney injury. Similar results were obtained in a study where bilateral IRI was induced in rats followed by either iv or renal artery administration of rat BM-MSCs transduced with a luciferase construct (Zhuo et al., 2013). The cells administered by either route appeared to localise to the lungs immediately after injection but had disappeared by day 7 after IRI.

Using CryoViz imaging, de Witte and colleagues had also demonstrated that human UC-MSCs are originally sequestered in the lungs and rapidly cleared after iv administration (de Witte et al., 2018). Furthermore, human A-MSCs infused iv after unilateral IRI in mice appeared only in the lungs over a 24-hour period (Luk et al., 2016). This is in contrast to a study by Wise and colleagues which reported that luciferase-labelled human BM-MSCs had reached the kidneys in mice with IRI and after immediate iv administration of cells (Wise et al., 2014), where they were detectable until day 3. In other studies from the same group using a unilateral ureteral obstruction mouse model, luciferase-labelled human BM-MSCs after iv administration were also reported in the kidneys (Huuskes et al., 2015; Wang et al., 2016). As far as we are aware, these three studies are the only ones to report the presence of luciferase-expressing cells in injured kidneys following iv administration. However, it is important to note that the cells in these studies were administered by the renal vein (Wise et al., 2014; Huuskes et al., 2015; Wang et al., 2016) which raises the possibility that the MSCs may have been delivered to the kidneys via retrograde flow.

## 6. CONCLUSIONS

In conclusion, using a combination of three experimental assessments as measures of kidney function and health in the same animal cohorts to determine therapeutic potential by direct comparison of three different MSC cell types in a mouse model of bilateral IRI revealed no convincing and consistently significant effect. We acknowledge the limited group sizes as a limitation of our study, preventing to draw robust conclusions. However, despite this limitation, our observations are in contrast to numerous studies reporting efficacy of either of the MSC types tested here but lacking direct side-by-side assessment in the same rodent model of AKI. Most published studies rely on serum biomarker and histology assessments and fail to incorporate longitudinal functional assessments. Further analyses will be required, in larger group sizes and including longitudinal measurements on kidney function, to determine whether factors including timing of administration and mouse strain/genetic background influence the therapeutic potential of MSCs.

## Supporting information

Supplementary Master Tables

## 7. LIST OF ABBREVIATIONS

AKI: acute kidney injury
A-MSC: adipose-derived mesenchymal stromal cell
BLI: bioluminescence imaging
BM-MSC: bone marrow-derived mesenchymal stromal cell
CKD: chronic kidney disease
ESKD: end stage kidney disease
GFR: glomerular filtration rate
IRI: ischaemia reperfusion injury
MSC: mesenchymal stromal cell
PBS: phosphate buffered saline
SD: standard deviation
UC-MSC: umbilical cord-derived mesenchymal stromal cell

## 8. DECLARATIONS

### Ethics approval

We obtained informed consent from all donors who kindly provided cells and tissue for our study. MSCs were obtained from different sites participating in the RenalToolBox network under respective MTAs with the University of Liverpool.

A-MSCs were provided by the University of Heidelberg (Germany) (title of approved project: ‘Isolation and characterisation of MSCs from human adipose tissue’; institutional approval committee: Mannheim Ethics Commission II; approval number: 2006-192N-MA; date of approval: 18.04.2005, re-confirmed 26.02.2009 with subsequent approvals).

BM-MSCs were provided by the University of Galway; cells had been isolated from bone marrow aspirates (healthy young male donors who had provided consent) purchased from Lonza (Basel, Switzerland).

UC-MSCs from two independent donors were obtained from the NHS Blood and Transplant and transferred to the University of Liverpool (project title: ‘The provision of mesenchymal stromal cells to the University of Liverpool for use in the RenalToolBox project’; institutional approval unit: NHS Blood and Transplant, Cellular and Molecular Therapies; approval number: RTB21112019; date of approval: 21 November 2019). All work was performed in line with the principles of the Declaration of Helsinki.

Ethical approval for the animal experiments was granted by the Home Office in accordance with UK regulations set out in the Animals (Scientific Procedures) Act 1986 and following approval by the Animal Welfare and Ethical Review Board (AWERB) of the University of Liverpool, under project licences PPL70_8741 (‘Preclinical therapies for renal and cardiovascular injury’, approved 6.10.2015) and PP3076489 (‘Developing safe and efficacious cell-based therapies for kidney disease’, approved 2.11.2020).

### Consent for publication

Not applicable

### Availability of data and materials

All datasets generated and analysed are contained within the article and the supplementary materials.

### Authors’ contributions

KTC and FA contributed to conception and design of the study, collection and assembly of data. LR contributed to the collection and analysis of data, data analysis and interpretation. MGF and DMH contributed to the experimental design and statistical analysis of the data, data analysis and interpretation. AT, VH, PM and BW contributed to conception and design, data analysis and interpretation. LR, MFG, AT, VH, PM and BW contributed to the supervision of the study. PM and BW obtained the funding of the study. All authors contributed to manuscript writing and final approval of the manuscript.

## Acknowledgements

We thank Sandra Calcat-i-Cervera (University of Galway) for providing the bone marrow-derived mesenchymal stromal cells, and Erika Erika and Eleonora Scaccia (University of Heidelberg) for providing the adipose-derived mesenchymal stromal cells.

We acknowledge expert support from the LIV-SRF Centre for Preclinical Imaging and the Biomedical Services Unit at the University of Liverpool for this study. We acknowledge technical support for the generation of tissue sections by the Liverpool University Biobank, and PAS staining by Veterinary Laboratory (Leahurst Campus), both University of Liverpool.

