## Supplementary Figures and Tables for "Assessing the efficacy of human mesenchymal stromal cells of different tissue origins in a mouse model of kidney ischaemia reperfusion injury"

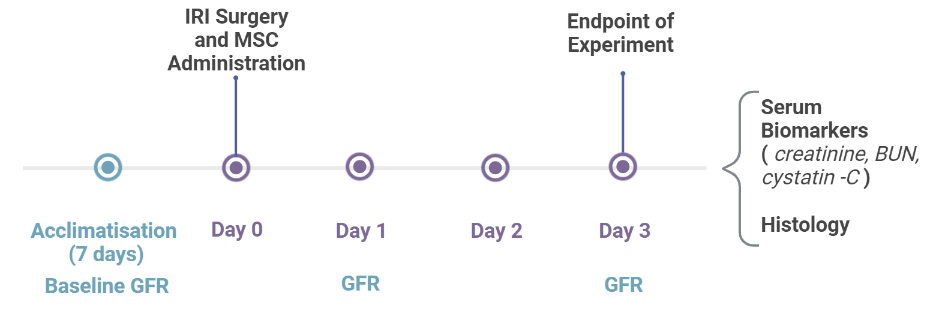

**Supplementary Figure 1.** **Experimental design to assess MSC efficacy in mice after IRI.** Transdermal GFR measurements were conducted at three different time points (baseline, day 1 and day 3 post-surgery). Cell administration was undertaken immediately after the IRI surgery and day 3 was the end point of the experiment.

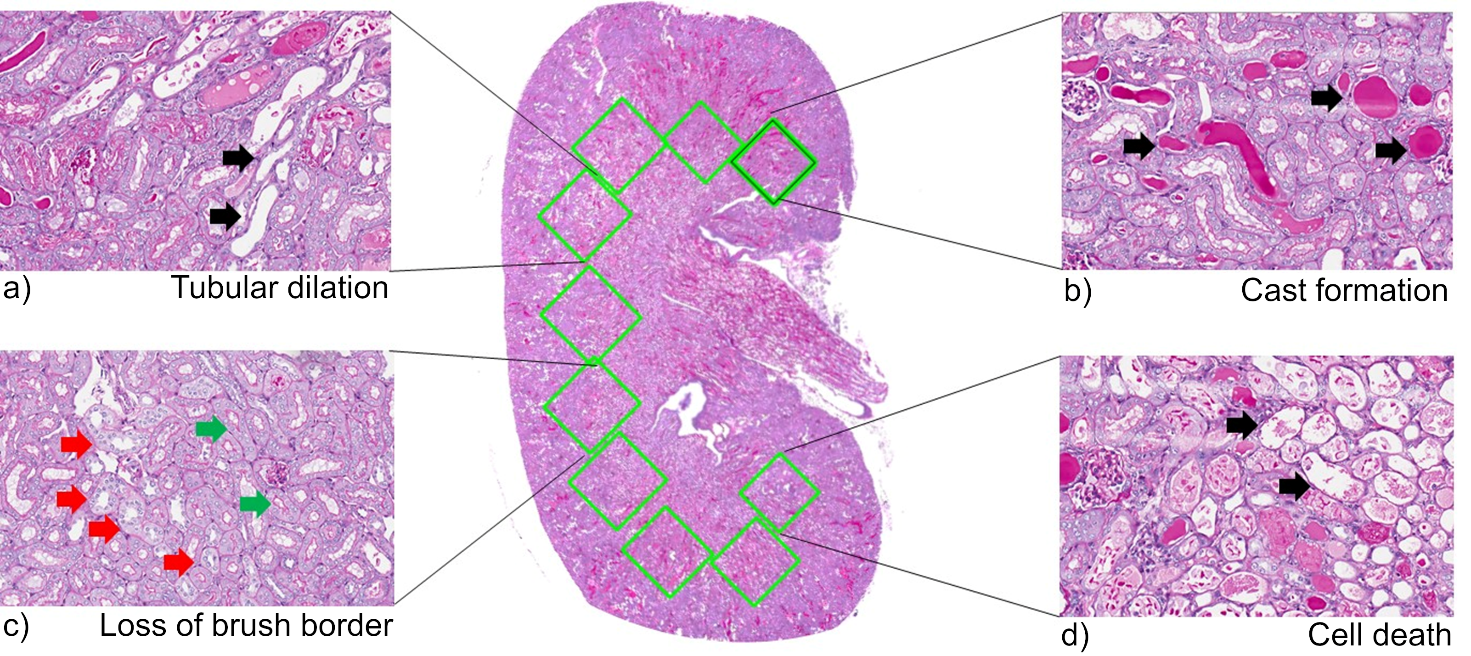

**Supplementary Figure 2.** **Histological changes of the outer stripe of the outer medulla (OSOM) in a mouse kidney after IRI.** Typical histopathological features of acute kidney injury were assessed by Periodic acid-Schiff (PAS). (**a**) Black arrows show the location of tubular dilation. (**b**) Black arrows indicate cast formation. (**c**) Green arrows point to the brush border of the proximal tubule in pink, while the red arrows show the loss of the tubular brush border. (**d**) Black arrows point out cell necrosis and fragmentation of the cytoplasm.

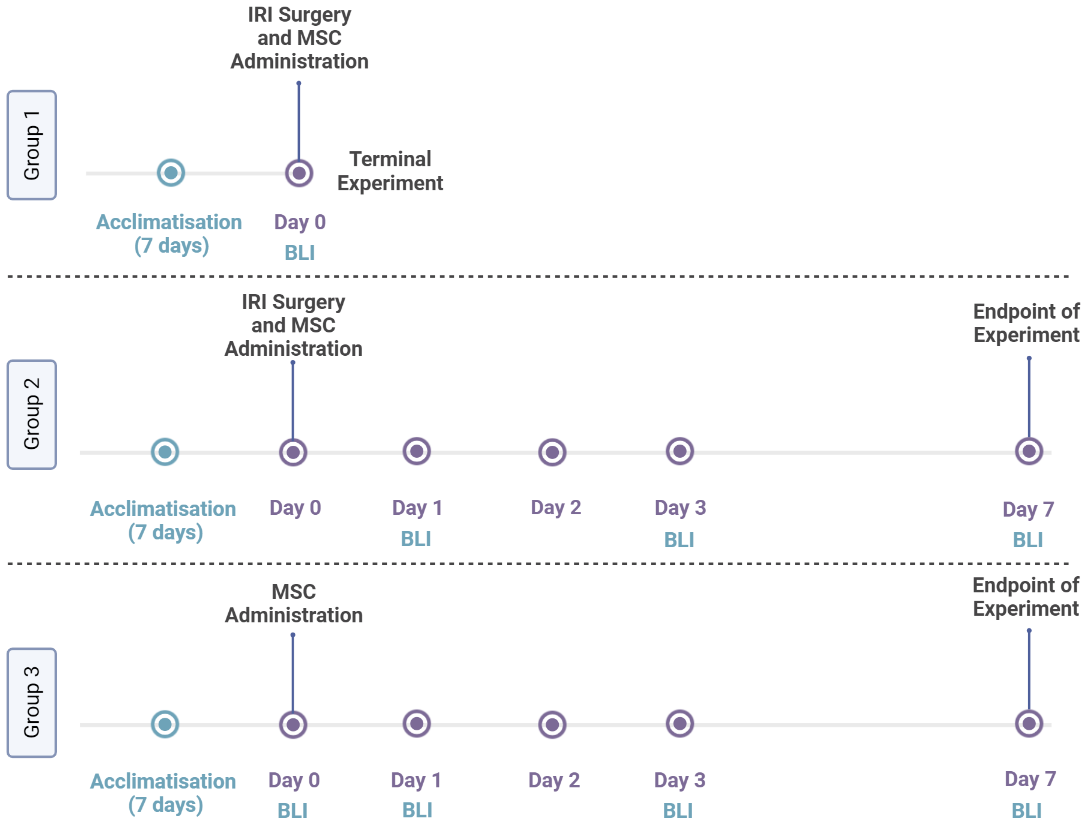

**Supplementary Figure 3.** **Experimental design to assess MSC biodistribution in mice after IRI.** Because IRI surgery, MSC administration and BLI imaging is too demanding on the animals’ wellbeing, BLI imaging on the day of the surgery was performed under terminal anaesthesia in group 1 animals (UC-MSC, n=4; BM-MSC, n=3; A-MSC, n=3; because UC-MSC administration was not complete in one animal, it was removed from the analysis). In group 2, mice underwent IRI surgery, then were allowed to recover after surgery and cell administration, and imaged on days 1, 3 and 7 (UC-MSC, n=5; BM-MSC, n=4; A-MSC, n=4; because IRI was too severe in one animal injected with UC-MSCs, it was terminated on day 1 and removed from the study). Mice in group 3 served as no-injury control group after receiving the cells but no IRI surgery, followed by imaging on days 1, 3 and 7. (UC-MSC, n=3; BM-MSC, n=3; A-MSC, n=3).

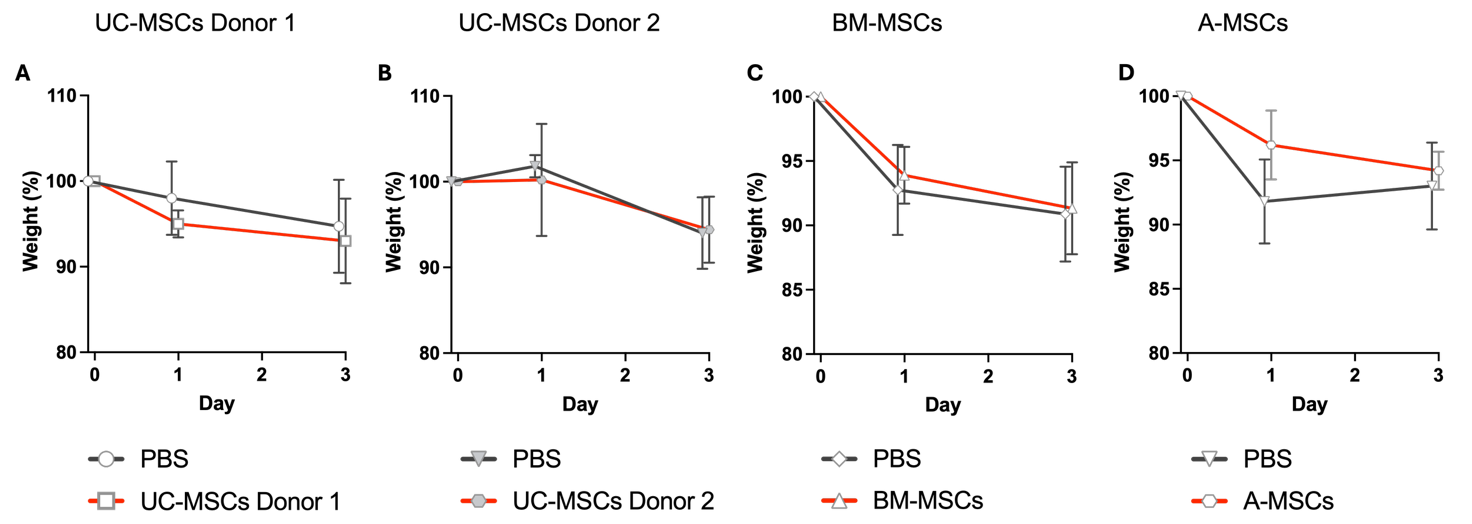

**Supplementary Figure 4. Changes in body weight in mice after renal IRI.** Comparison between controls and groups treated with **(A)** UC-MSCs from donor 1, **(B)** UC-MSCs from donor 2, **(C)** BM-MSCs and **(D)** A-MSCs. Data are from animals of the efficacy study (SI Master Table I). Data shown as mean $\pm$ SD.

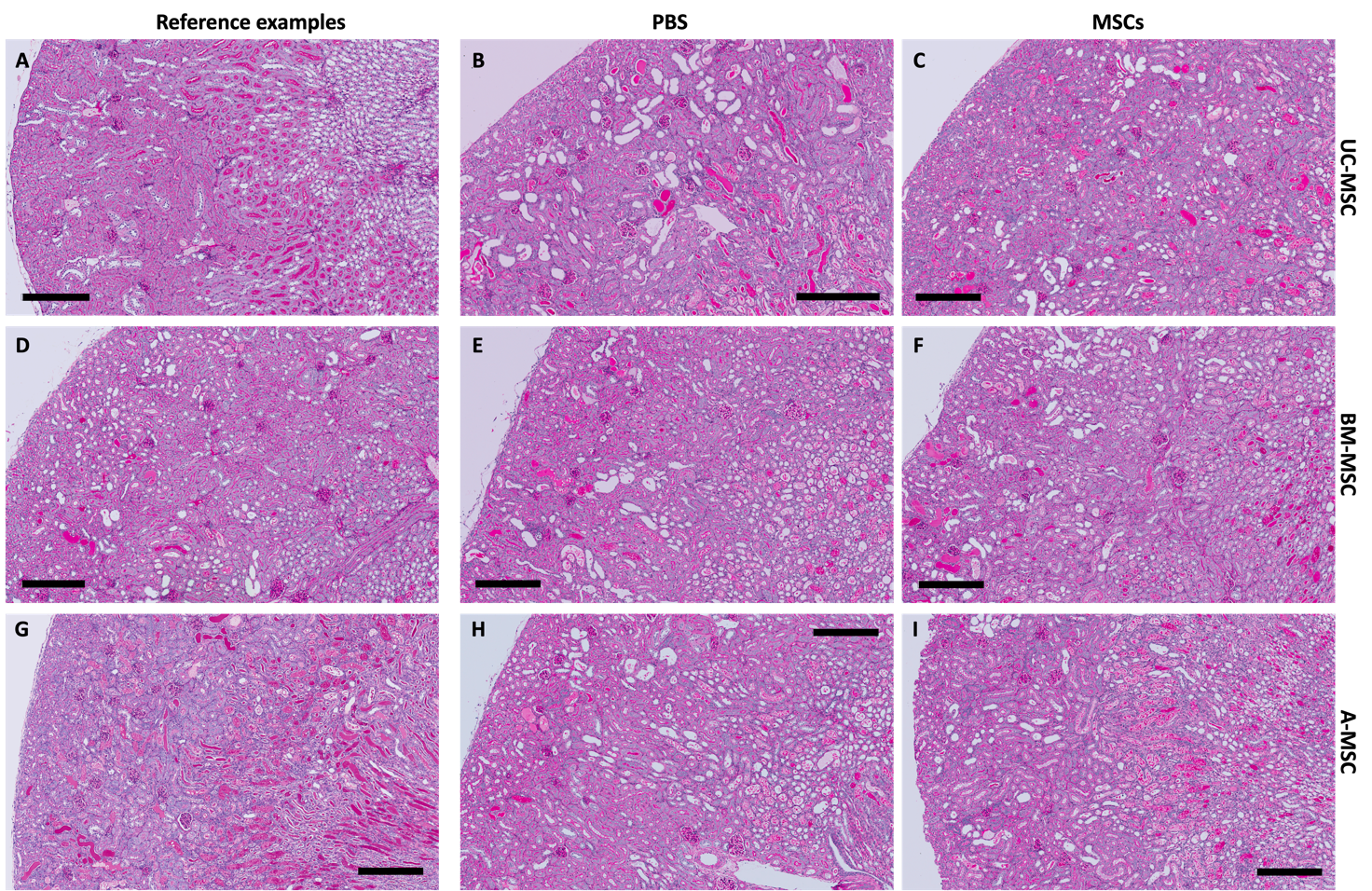

**Supplementary Figure 5. Histological analysis of kidney sections stained with Periodic acid-Schiff (PAS). (A)** Representative image of healthy tissue from uninjured animal. **(B)** Representative image of injured tissue with < 10% (Wang score 1.10) damage. **(G)** Representative image of injured tissue with > 75% damage (Wang score 3.75) **(B, E, H)** Representative images of kidney sections from IRI-injured, PBS-treated control groups, and **(C, F, I)** from IRI-injured, cell-treated groups (MSCs) on day 3 after IRI. **(B, C)** show images of UC-MSC cohort, **(E, F)** of the BM-MSC cohort, and **(H, I)** of the A-MSC cohort). Scale bars, 300μm.

**Supplementary Table 1.** **Tabulated values of the mean GFR for each experimental group.** Data shown as mean and standard deviation (SD).

| **Group** | **GFR (µL/(min * 100 g bw))** | | | | | |
| --- | --- | --- | --- | --- | --- | --- |
|  | **Day 0** | | **Day 1** | | **Day 3** | |
|  | **Mean** | **SD** | **Mean** | **SD** | **Mean** | **SD** |
| **IRI + PBS (n=8)** | 1278.4 | 109.0 | 362.0 | 152.6 | 701.6* | 239.9* |
| **IRI + UC-MSC Donor 1 (n=6)** | 1293.2 | 194.3 | 291.0 | 199.6 | 677.4 | 144.9 |
| **IRI + PBS (n=7)** | 1276.9 | 204.9 | 329.9 | 187.6 | 715.9 | 309.2 |
| **IRI + UC-MSC Donor 2 (n=6)** | 1177.0 | 302.1 | 292.4** | 208.4** | 490.1 | 395.5 |
| **IRI + PBS (n=8)** | 1198.3 | 179.1 | 242.7 | 161.9 | 670.3 | 292.1 |
| **IRI + BM-MSC (n=9)** | 1190.4 | 139.2 | 364.3 | 189.9 | 843.8 | 264.6 |
| **IRI + PBS (n=10)** | 1194.7 | 182.61 | 347.3 | 169.2 | 839.5 | 257.6 |
| **IRI + A-MSC (n=9)** | 1200.9 | 141.3 | 411.1 | 344.4 | 1053.3 | 388.6 |

Min = minute; bw = bodyweight (in gram, g); * - data from 7 animals; ** - data from 4 animals.

**Supplementary Table 2.** **Changes in kidney function as percentage GFR for each experimental group.** Percentage of kidney function relative to baseline values. Data shown as mean and standard deviation (SD).

| **Group** | **Percentage kidney function based on GFR (%)** | | | |
| --- | --- | --- | --- | --- |
|  | **Day 1** | | **Day 3** | |
|  | **Mean** | **SD** | **Mean** | **SD** |
| **IRI + PBS (n=8)** | 28.8 | 13.1 | 56.6* | 21.4* |
| **IRI + UC-MSCs Donor 1 (n=6)** | 22.4 | 13.9 | 54.0 | 17.5 |
| **IRI + PBS (n=7)** | 26.5 | 14.3 | 58.0 | 27.8 |
| **IRI + UC-MSCs Donor 2 (n=6)** | 21.8** | 17.1** | 38.8 | 29.8 |
| **IRI + PBS (n=8)** | 20.3 | 11.7 | 56.7 | 23.3 |
| **IRI + BM-MSCs (n=9)** | 30.5 | 15.9 | 71.0 | 21.2 |
| **IRI + PBS (n=10)** | 29.1 | 14.1 | 69.8 | 16.4 |
| **IRI + A-MSCs (n=9)** | 33.8 | 27.0 | 88.9 | 33.8 |

* - data from 7 animals; ** - data from 4 animals.

**Supplementary Table 3: Linear Mixed Model Coefficients for GFR change over time.**

|  |  | **Estimate** | **Std. Error** | **95% Confidence Interval** | **p-value** |
| --- | --- | --- | --- | --- | --- |
| **PBS**  **Reference group** | **Intercept** | 1305.4 | 67.5 | (1180, 1430.7) | **<0.001** |
|  | **Slope1 for time (0, 1)** | -863.7 | 45.8 | (-950.5, -776.9) | **<0.001** |
|  | **Slope2 for time (1, 3)** | 238.3 | 23.1 | (194.4, 282.1) | **<0.001** |
| **UC-MSCs (Donor 1)**  **vs PBS** | **Diff Constant** | 86.6 | 121.2 | (-138.8, 312.1) | 0.48 |
|  | **Diff Slope1 for time (0, 1)** | -138.4 | 116.8 | (-359.7, 82.9) | 0.24 |
|  | **Diff Slope2 for time (1, 3)** | -45.1 | 58.5 | (-155.9, 65.8) | 0.44 |
| **UC-MSCs (Donor 2)**  **vs PBS** | **Diff Constant** | -14.1 | 123.6 | (-244, 215.6) | 0.91 |
|  | **Diff Slope1 for time (0, 1)** | -108.1 | 131.7 | (-353.8, 147.4) | 0.41 |
|  | **Diff Slope2 for time (1, 3)** | -95.9 | 66.0 | (-223.6, 27.2) | 0.15 |
| **BM-MSCs**  **vs PBS** | **Diff Constant** | -18.1 | 108.5 | (-219.9, 183.6) | 0.87 |
|  | **Diff Slope1 for time (0, 1)** | -91.9 | 103.7 | (-288.4, 104.5) | 0.38 |
|  | **Diff Slope2 for time (1, 3)** | -24.5 | 52.0 | (-122.8, 74) | 0.64 |
| **A-MSCs**  **vs PBS** | **Diff Constant** | -110.7 | 101.6 | (-299.7, 78.2) | 0.28 |
|  | **Diff Slope1 for time (0, 1)** | 16.4 | 95.0 | (-163.6, 196.4) | 0.86 |
|  | **Diff Slope2 for time (1, 3)** | 7.8 | 47.6 | (-82.3, 98.1) | 0.87 |
| **PBS heterogeneity (Reference experiment: A-MSCs-PBS)** | **Experiment UC-MSCs vs PBS** | -98.9 | 91.0 | (-267.7, 70) | 0.28 |
|  | **Experiment BM-MSCs vs PBS** | -88.9 | 87.7 | (-251.9, 74.1) | 0.32 |
|  | **Experiment UC-MSCs (Donor 2) vs PBS** | -114.2 | 93.8 | (-288.5, 60.1) | 0.23 |

**Explanation:** time (0, 1) refers to the slope for the time between d0 and d1; time (1, 3) refers to the slope for the time between d1 and d3.

**Supplementary Table 4.** **Tabulated values for the biomarker measurements at Day 3.** Data shown as mean and standard deviation (SD).

| **Group** | **Creatinine (mg/dL)** | | **BUN (mg/dL)** | | **Cystatin-C (mg/L)** | |
| --- | --- | --- | --- | --- | --- | --- |
|  | **Mean** | **95% CI** | **Mean** | **95% CI** | **Mean** | **95% CI** |
| **Healthy reference animals (uninjured)** | 0.64 * | 0.56 – 0.73 | 28.43 ** | 20.16 – 36.70 | 0.39 *** | 0.33 – 0.45 |
|  | **Mean** | **SD** | **Mean** | **SD** | **Mean** | **SD** |
| **IRI + PBS** (n=8) | 0.90 | 0.14 | 67.10 | 25.50 | 0.71^$^ | 0.08^$^ |
| **IRI + UC-MSCs** (n=5) | 0.78 | 0.13 | 62.11 | 26.17 | 0.75^ | 0.21^ |
| **IRI + PBS** (n=8) | 0.78 | 0.08 | 71.96 | 37.11 | 0.69 | 0.16 |
| **IRI + BM-MSCs** (n=9) | 0.61 | 0.14 | 63.66 | 24.38 | 0.59 | 0.13 |
| **IRI + PBS** (n=10) | 0.69 | 0.10 | 64.17 | 24.61 | 0.63 | 0.14 |
| **IRI + A-MSCs** (n=9) | 0.71 | 0.14 | 58.51 | 24.91 | 0.62 | 0.18 |

* n = 10 animals; ** n = 6 animals; *** n = 7 animals.

^$^ Only 7 samples measured for this specific biomarker and group.

^ Only 4 samples measured for this specific biomarker and group.

For details, refer to Supplementary Master Table I and III.

**Supplementary Table 5: Linear regression models for serum biomarkers at Day 3.**

|  |  | **Estimate** | **Std. Error** | **95% Confidence Interval** | **p-value** |
| --- | --- | --- | --- | --- | --- |
| ***Creatinine (mg/dL)*** | | | | | |
|  | **Intercept** | 0.69 | 0.04 | (0.62, 0.77) | <0.001 |
| **Treatment (Reference = PBS)** | **A-MSCs** | 0.02 | 0.06 | (-0.09, 0.13) | 0.73 |
|  | **BM-MSCs** | -0.18 | 0.06 | (-0.29, -0.06) | **0.005** |
|  | **UC-MSCs** | -0.13 | 0.07 | (-0.27, 0.01) | 0.08 |
| **PBS heterogeneity (Reference experiment: A-MSCs-PBS)** | **Experiment: BM-MSCs vs PBS** | 0.09 | 0.06 | (-0.03, 0.2) | 0.14 |
|  | **Experiment: UC-MSCs vs PBS** | 0.21 | 0.06 | (0.09, 0.33) | <0.001 |
| ***Cystatin-C (mg/L)*** | | | | | |
|  | **Intercept** | 0.63 | 0.05 | (0.53, 0.72) | <0.001 |
| **Treatment (Reference = PBS)** | **A-MSCs (vs PBS)** | -0.01 | 0.07 | (-0.15, 0.13) | 0.65 |
|  | **BM-MSCs (vs PBS)** | -0.10 | 0.07 | (-0.25, 0.05) | 0.17 |
|  | **UC-MSCs (vs PBS)** | 0.09 | 0.09 | (-0.1, 0.28) | 0.86 |
| **PBS heterogeneity (Reference experiment: A-MSCs-PBS)** | **Experiment: BM-MSCs vs PBS** | 0.06 | 0.07 | (-0.08, 0.2) | 0.41 |
|  | **Experiment: UC-MSCs vs PBS** | 0.08 | 0.07 | (-0.07, 0.23) | 0.28 |
| ***Log(Bun (mg/DL))*** | | | | | |
|  | **Intercept** | 4.09 | 0.13 | (3.83, 4.35) | <0.001 |
| **Treatment (Reference = PBS)** | **A-MSCs (vs PBS)** | -0.11 | 0.19 | (-0.49, 0.26) | 0.55 |
|  | **BM-MSCs (vs PBS)** | -0.08 | 0.20 | (-0.48, 0.32) | 0.70 |
|  | **UC-MSCs (vs PBS)** | 0.00 | 0.23 | (-0.47, 0.47) | 0.99 |
| **PBS heterogeneity (Reference experiment: A-MSCs-PBS)** | **Experiment: BM-MSCs vs PBS** | 0.07 | 0.19 | (-0.31, 0.46) | 0.70 |
|  | **Experiment: UC-MSCs vs PBS** | 0.06 | 0.19 | (-0.33, 0.45) | 0.77 |

**Supplementary Table 6.** **Tabulated values for the histological analysis (Wang score).** Data shown as mean and standard deviation (SD).

**(A)	(B)**

| **Combined Wang Scores, PBS vs MSC** | | |
| --- | --- | --- |
| **Group** | **Wang Score** | |
|  | **Mean** | **SD** |
| **IRI + PBS (n=8)** | 2.47 | 0.55 |
| **IRI + UC-MSCs (n=6)** | 2.38 | 0.59 |
| **IRI + PBS (n=4)** | 2.55 | 0.65 |
| **IRI + BM-MSCs (n=4)** | 2.34 | 0.58 |
| **IRI + PBS (n=5)** | 2.48 | 0.69 |
| **IRI + A-MSCs (n=4)** | 2.60 | 0.29 |

| **Left vs Right Score, all kidneys** | | | |
| --- | --- | --- | --- |
| **Group** | **Wang Score** | | |
|  | **Mean** | **SD** | **p-value** |
| **Left Kidneys (n=31)** | 2.13 | 0.65 | **0.0001** |
| **Right Kidneys (n=29)** | 2.84 | 0.69 |  |

**(C)**

| **Left vs Right Score, UC-MSCs** | | | |
| --- | --- | --- | --- |
| **Group** | **Wang Score** | | |
|  | **Mean** | **SD** | **p-value** |
| **IRI+PBS, Left (n=8)** | 1.94 | 0.73 | 0.12 |
| **IRI+UC-MSC, Left (n=6)** | 2.48 | 0.49 |  |
| **IRI+PBS, Right (n=7)** | 3.14 | 0.58 | 0.09 |
| **IRI+UC-MSC, Right (n=5)** | 2.25 | 0.88 |  |

**(D)**

| **Left vs Right Score, BM-MSCs** | | | |
| --- | --- | --- | --- |
| **Group** | **Wang Score** | | |
|  | **Mean** | **SD** | **p-value** |
| **IRI+PBS, Left (n=4)** | 2.53 | 0.80 | 0.14 |
| **IRI+BM-MSC, Left (n=4)** | 1.68 | 0.56 |  |
| **IRI+PBS, Right (n=4)** | 2.58 | 0.58 | 0.36 |
| **IRI+BM-MSC, Right (n=4)** | 3.00 | 0.62 |  |

**(E)**

| **Left vs Right Score, A-MSCs** | | | |
| --- | --- | --- | --- |
| **Group** | **Wang Score** | | |
|  | **Mean** | **SD** | **p-value** |
| **IRI+PBS, Left (n=5)** | 2.15 | 0.79 | 0.71 |
| **IRI+A-MSC, Left (n=4)** | 2.01 | 0.23 |  |
| **IRI+PBS, Right (n=5)** | 2.81 | 0.70 | 0.39 |
| **IRI+A-MSC, Right (n=4)** | 3.18 | 0.41 |  |

**Supplementary table 7: Linear regression models for Mean Wang score**

|  |  | **Estimate** | **Std. Error** | **95% Confidence Interval** | **P-value** |
| --- | --- | --- | --- | --- | --- |
|  | **Intercept** | 2.484 | 0.257954 | (1.95, 3.02) | <0.001 |
| **Treatment (Reference = PBS)** | **A-MSCs (vs PBS)** | 0.111 | 0.386931 | (-0.69, 0.91) | 0.78 |
|  | **BM-MSCs (vs PBS)** | -0.2075 | 0.407861 | (-1.05, 0.63) | 0.62 |
|  | **UC-MSCs (vs PBS)** | -0.09083 | 0.311509 | (-0.73, 0.55) | 0.77 |
| **PBS heterogeneity (Reference experiment: A-MSCs-PBS)** | **Experiment:**  **BM-MSCs vs PBS** | 0.066 | 0.386931 | (-0.73, 0.86) | 0.87 |
|  | **Experiment:**  **UC-MSCs vs PBS** | -0.0115 | 0.328828 | (-0.69, 0.67) | 0.97 |

**Supplementary table 8.** **Tabulated values for the bioluminescence measurements.** Data shown as mean values and standard deviation (SD) [p/s/cm^2^/sr].

| **Group** | **Day 0** | | **Day 1** | | **Day 3** | | **Day 7** | |
| --- | --- | --- | --- | --- | --- | --- | --- | --- |
|  | **Mean** | **SD** | **Mean** | **SD** | **Mean** | **SD** | **Mean** | **SD** |
| **IRI + UC-MSCs (n=3) (group 1)** | 7.3x10^7^ | 1.3x10^7^ |  |  |  |  |  |  |
| **IRI + UC-MSCs (n=4) (group 2)** |  |  | 1.3x10^7^ | 0.5x10^7^ | 5.8x10^5^ | 3.2x10^5^ | 1.6x10^5^ | 0.2x10^5^ |
| **Healthy controls + UC-MSCs (n=3) (group 3)** | 7.5x10^7^ | 1.3x10^7^ | 3.8x10^6^ | 0.8x10^6^ | 4.2x10^5^ | 1.0x10^5^ | 1.3x10^5^ | 0.1x10^5^ |
| **IRI + BC-MSCs (n=3)**  **(group 1)** | 8.5x10^7^ | 0.5x10^7^ |  |  |  |  |  |  |
| **IRI + BC-MSCs (n=4)**  **(group 2)** |  |  | 6.5x10^6^ | 3.2x10^6^ | 2.3x10^5^ | 0.89x10^5^ | 1.3x10^5^ | 0.2x10^5^ |
| **Healthy controls + BC-MSCs (n=3) (group 3)** | 4.8x10^7^ | 1.0x10^7^ | 1.4x10^6^ | 0.6x10^6^ | 2.0x10^5^ | 0.16x10^5^ | 1.1x10^5^ | 0.1x10^5^ |
| **IRI + A-MSCs (n=3)**  **(group 1)** | 5.9x10^7^ | 2.4x10^7^ |  |  |  |  |  |  |
| **IRI + A-MSCs (n=4)**  **(group 2)** |  |  | 3.0x10^7^ | 1.0x10^7^ | 3.0x10^6^ | 1.6x10^6^ | 1.8x10^5^ | 0.5x10^5^ |
| **Healthy controls + A-MSCs (n=3) (group 3)** | 3.3x10^7^ | 1.4x10^7^ | 4.0x10^6^ | 1.3x10^6^ | 1.1x10^6^ | 0.6x10^6^ | 1.2x10^5^ | 0.1x10^5^ |
